# Systematic functional annotation workflow for insects

**DOI:** 10.1101/2022.05.12.490705

**Authors:** Hidemasa Bono, Takuma Sakamoto, Takeya Kasukawa, Hiroko Tabunoki

**Affiliations:** Laboratory of Bio-DX, Genome Editing Innovation Center, Hiroshima University, 3-10-23 Kagamiyama, Higashi-Hiroshima city, Hiroshima, 739-0046, Japan; Laboratory of Genome Informatics, Graduate School of Integrated Sciences for Life, Hiroshima University, 3-10-23 Kagamiyama, Higashi-Hiroshima city, Hiroshima, 739-0046, Japan; Institute of Global Innovation Research, Tokyo University of Agriculture and Technology, 3-5-8 Saiwai-cho, Fuchu, Tokyo, 183-8509; Department of Science of Biological Production, Graduate School of Agriculture, Tokyo University of Agriculture and Technology, 3-5-8 Saiwai-cho, Fuchu, Tokyo, 183-8509, Japan; RIKEN Center for Integrative Medical Sciences, 1-7-22 Suehiro-cho, Tsurumi-ku, Yokohama, Kanagawa, 230-0045, Japan

**Keywords:** Functional annotation, RNA sequencing, Transcriptome assembly, Stick insect, Silkworm

## Abstract

Next generation sequencing has revolutionized entomological study, rendering it possible to analyze the genomes and transcriptomes of non-model insects. However, use of this technology is often limited to obtaining nucleotide sequences of target or related genes, with many of the acquired sequences remaining unused because other available sequences are not sufficiently annotated. To address this issue, we have developed a functional annotation workflow for transcriptome-sequenced insects to determine transcript descriptions, which represents a significant improvement over the previous method (functional annotation pipeline for insects). The developed workflow attempts to annotate not only the protein sequences obtained from transcriptome analysis but also the ncRNA sequences obtained simultaneously. In addition, the workflow integrates the expression level information obtained from transcriptome sequencing for application as functional annotation information. Using the workflow, functional annotation was performed on the sequences obtained from transcriptome sequencing of stick insect (*Entoria okinawaensis*) and silkworm (*Bombyx mori*), yielding richer functional annotation information than that obtained in our previous study. The improved workflow allows more comprehensive exploitation of transcriptome data and is applicable to other insects because the workflow has been openly developed on GitHub.

**Simple Summary:** The function of all genes encoded in the genome should be studied for genome editing. The genome editing technology can speeds up insect research for functional analysis of genes. Our knowledge about the functional information of genes is still incomplete currently while genome sequencing of an organism can be completed. The functional information has been annotated based solely on the information that has been obtained from the result of previous biological research. However, this information will be important in determining the target genes for genome editing. In particular, it is very important that this information is in machine-readable form because computer programs mainly parse this information for the understanding of biological systems. In this paper, we describe a workflow-based method for annotating gene functions in insects that make use of transcribed sequence information as well as reference genome and protein sequence databases. Using the developed workflow, we annotated functional information of Japanese stick insect and silkworm, including gene expression as well as sequence analysis. The functional annotation information obtained by the workflow will greatly expand the possibilities of entomological research using genome editing.

## 1. Introduction

Genome sequencing is becoming common practice in insect research. As of May 2022, the genomes of around 3,000 insect species have been decoded and registered in the genome section of NCBI Datasets [1]. In addition, long-read technology is further accelerating the pace of insect genome sequencing [2].

However, some insects have genomes larger than that of humans, further complicating the difficult process of whole genome sequencing. As an alternative, transcriptome sequencing using next generation sequencing technology, also termed RNA sequencing (RNA-Seq), provides a powerful tool for evaluating non-model species such as large-genome-size insects [3]. In particular, this strategy can efficiently identify tens of thousands of possible genes in a specific tissue by assembling tens of millions of reads. The sequences are then assembled for the identification of transcriptional units. Systematic sequence similarity searches against non-redundant sequence sets and BLAST2GO searches [4] are often employed for the functional inference of such transcriptomes to be utilized in gene set enrichment analyses. Moreover, a new method termed Seq2Fun that does not require transcriptome *de novo* assembly was recently developed for rapid functional profiling of RNA-Seq data for non-model organisms [5]. Nevertheless, the output of such analyses is reliant upon the comprehensiveness of the available datasets and their functional annotation.

Traditionally, reference data have been maintained as research community databases (DBs) such as FlyBase for fruit flies [6]. In turn, such communities spearheaded the development of Gene Ontology (GO) [7], which provides machine-interpretable annotations incorporating human-readable descriptions. These annotations for genomic research, including GO, are provided by Ensembl and cover a variety of vertebrates [8]. For insects, Ensembl Metazoa in Ensembl Genomes integrates the community annotations of individual insects [9]. Nevertheless, it is unable to keep pace with the increase of insect genome sequencing including currently available genome and transcriptome data.

Concurrently, we developed a functional annotation pipeline in non-model insects for microarray-based transcriptome analysis of a unique mutant of the silkworm *Bombyx mori op* [10]. Subsequently, we also utilized this pipeline for comparative studies with humans and flies [11, 12]. In particular, by assigning human gene identifiers (IDs) to the silkworm genes, we could facilitate the reconstruction of pathway DBs, which were originally developed for the analysis of model organisms [10,13]. However, such prior studies have almost exclusively been limited to protein coding sequences, as functional annotation pipelines for transcriptome sequences were originally designed for mammalian transcriptomes [14] produced by sequencing cDNAs including expressed sequence tags (ESTs) using Sanger sequencers in the Functional Annotation of Mammalian genomes (FANTOM) project [15, 16]. Although functional annotation for non-coding RNAs (ncRNA) was considered in the pipeline for FANTOM3 [17], this feature was ultimately not incorporated because the reference information and biological knowledge regarding ncRNAs were not yet sufficient [18].

As transcriptome sequencing becomes popular, many groups run pipelines of their own, with the information regarding the transcription units from various studies being reported on a study-by-study basis. Moreover, the transcript assemblies are often not registered in a public DB even though it is recommended that such data be deposited in the Transcriptome Shotgun Assembly (TSA) sequence database under the International Nucleotide Sequence Database Collaboration (INSDC) [19, 20], as the data need to be annotated uniformly to facilitate reuse in other analyses. However, the functional information of the currently registered transcription units is not sufficiently rich to support functional analysis and almost never annotated once archived even if it is available. Thus, the functional annotation of transcriptomes in a manner that can be integrated with datasets from other groups including the data in public DBs is urgently required.

Therefore, in this study we evaluated the functional annotation of transcripts from assembled transcriptomes to establish a functional annotation workflow for insect transcriptomes by updating the previously developed functional annotation pipeline for silkworm [10]. Then, we developed a systematic functional annotation workflow for insect research termed “Fanflow4Insects”. Fanflow4Insects enables the automatic annotation of sequences to exploit the exponential increase in genome and transcriptome information. Finally, Fanflow4Insects was tested using not only the reference transcriptome of silkworm (*Bombyx mori*) but also transcriptomes of the Japanese stick insect (*Entoria okinawaensis*) including the newly sequenced transcriptome of the fat body.

## 2. Materials and Methods

### 2.1. Programs comprizing the workflow

All scripts that form the workflow described below are freely accessible from the Systematic Analysis for Quantification of Everything (SAQE) [21] code repository at GitHub [22] along with the detailed parameters.

For transcriptomes without reference genome sequence, we utilized Trinity (v2.12.0) with paired-end parameters in the Docker environment (trinityrnaseq/trinityrnaseq:2.12.0) to construct *de novo* transcriptome assemblies [23].

Protein coding sequences within transcriptome sequences produced by *de novo* assembly or by mapping the reference genome sequence were predicted using TransDecoder (v5.2.0) [24]. The predicted and then translated protein sequences were subsequently compared via successive execution of the GGSEARCH program (ggsearch36; v36.3.8g) with ‘-d1 -m 10 -E 0.1’ parameters in the FASTA package [25] against the protein reference sequence DBs described in Table 1. The protein domain search program (hmm-scan) in the HMMER package [26] was used for the predicted protein sequences. Pfam (v35.0) was used as the protein domain DB [27].

**Table 1.** Source of reference databases.

| Category | Name of resource | URL |
| --- | --- | --- |
| Protein sequences | Ensembl | <sup>1</sup> https://ftp.ensembl.org/pub/release-105/fasta/homo_sapiens/pep/Homo_sapiens.GRCh38.pep.all.fa.gz |
|  | UniProtKB | https://ftp.uniprot.org/pub/databases/uniprot/current_release/knowledge-base/complete/uniprot_sprot.fasta.gz |
| Non-coding RNA sequences | Ensembl | <sup>1</sup> http://ftp.ensembl.org/pub/release-105/fasta/homo_sapiens/ncrna/Homo_sapiens.GRCh38.ncrna.fa.gz |
|  | EnsemblGenomes | <sup>1</sup> http://ftp.ensemblgenomes.org/pub/release-52/metazoa/fasta/drosophila_melanogaster/ncrna/Drosophila_melanogaster.BDGP6.32.ncrna.fa.gz |
| Protein and RNA domain | Pfam | http://ftp.ebi.ac.uk/pub/databases/Pfam/releases/Pfam35.0/Pfam-A.hmm.gz |
|  | Rfam | http://ftp.ebi.ac.uk/pub/databases/Rfam/14.7/Rfam.cm.gz |

<sup>1</sup> Only the URL for reference data of the typical organism is listed.

The transcriptome sequences were then used for annotation as ncRNA sequences. Nucleotide BLAST (blastn; v2.6.0+) [28] was used to detect sequence similarity to known ncRNA sequences in Ensembl (human) [8] and Ensembl Genomes (fly) [9] DBs. The program (cmscan) in the Infernal: inference of the RNA alignments package (v1.1.4) [29], which was developed for searching nucleotide sequence DBs for RNA structure and sequence similarities, was used to annotate transcript sequences against Rfam (v14.7), utilized as the RNA family DB [30].

A program (align_and_estimate_abundance.pl) in the Docker container for Trinity (trinityrnaseq/trinityrnaseq:2.11.0) was used to estimate the abundance of RNA-Seq reads [23]. Salmon (v1.0.0) [31] was selected to quantify the transcripts to produce the expression matrix for each organism.

### 2.2. Insects and sample collection

The stick insects *E. okinawaensis* were obtained from Amami-Ohshima in Kagoshima, Japan, in 2011 and Ishigaki Island in Okinawa, Japan, in 2013, as described in our previous study [32]. Three biological replicates of the fat body were dissected from adult *E. okinawaensis*. These tissues were stored at −80 °C until use. The fat body samples were weighed, homogenized with lysis buffer from a PureLink® RNA extraction kit (Thermo Fisher Scientific Inc., Valencia, CA, USA), and then centrifuged at 13,000×*g* for 10 min. The supernatants were then collected and processed for RNA purification according to the manufacturer’s instructions. Purified total RNA samples (1 µg each) were processed for RNA sequencing.

### 2.3. RNA sequencing

RNA quality was assessed using a Bioanalyzer 2100 (Agilent Technologies, Santa Clara, CA, USA). Libraries for cDNA sequencing were constructed using the Illumina TruSeq v2 kit (Illumina Inc., San Diego, CA, USA) according to the manufacturer’s protocol, and 100 bp paired-end sequencing was then performed on the HiSeq 2500 platform (Illumina). Using Trim Galore! (v0.4.4;) [33], low-quality bases and adapter sequences were trimmed, and the data quality was visualized and then confirmed. The raw RNA sequence reads for biological triplicates of *E. okinawaensis* fat body samples were deposited in the Sequence Read Archive (SRA) under accession ID: DRA013458 (DRR346171, DRR346172, and DRR346173).

## 3. Results

### 3.1. Functional annotation workflow for insects

We first evaluated rules for the functional annotation of transcripts. In our previous studies, we only focused on translated protein sequences [12,32]. In the present study, we also aimed to annotate sequences without coding regions using ncRNA references, and quantified expression information from RNA-Seq as well as protein coding sequences ascertained based on sequence similarity. For transcripts with no sequence similarity, quantified expression values were used for annotation (Figure 1). The functional annotation workflow described below, being developed especially for insects, was termed “Fan-flow4Insects”.

**Figure 1.**
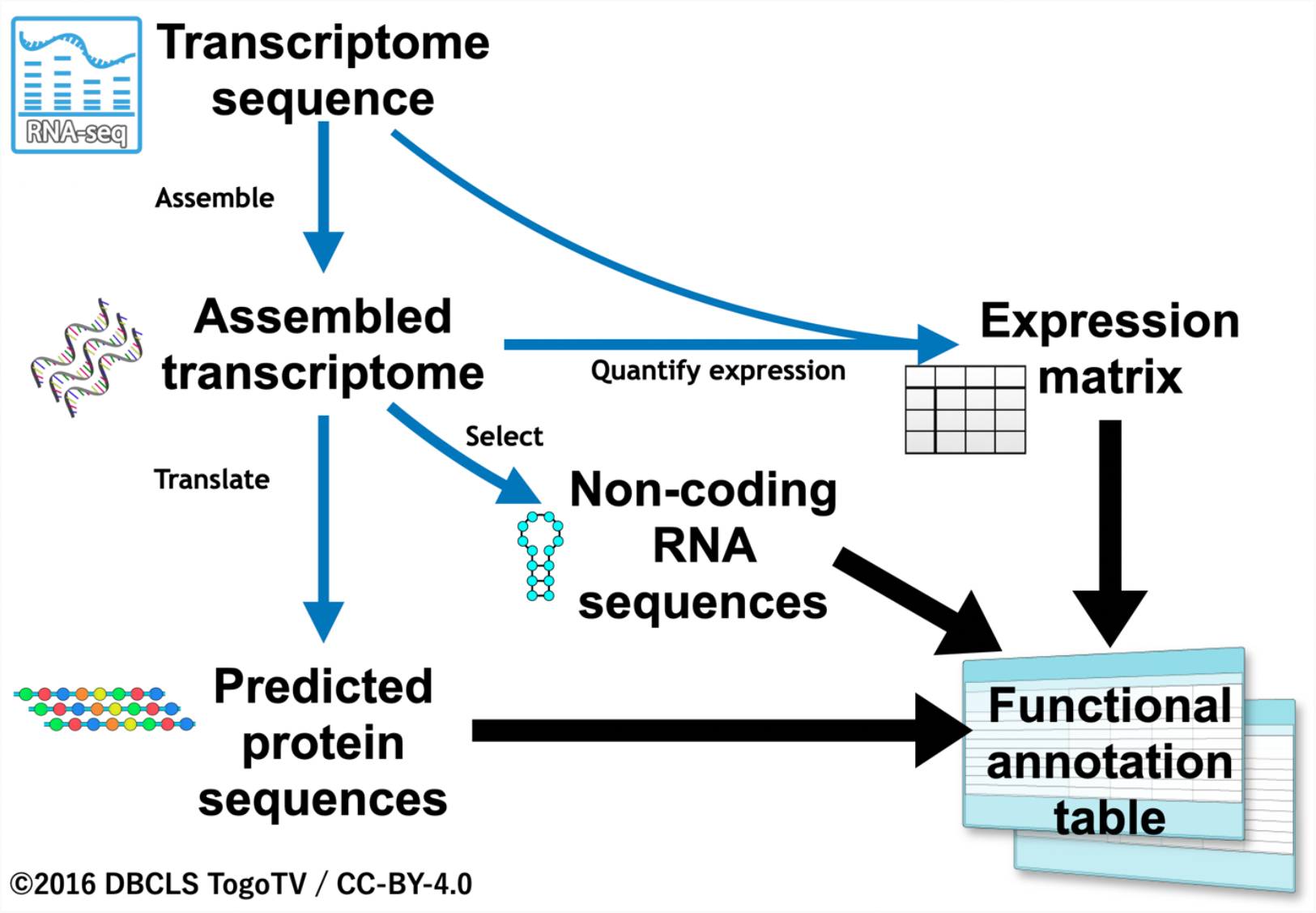
Overview of the annotation workflow, Fanflow4Insects.

#### 3.1.1. Functional annotation of coding sequences

Using predicted protein sequences translated from each assembled transcriptome, we performed systematic sequence similarity searches against the predicted protein sequence datasets of functionally well-annotated organisms, including human (*Homo sapiens*), mouse (*Mus musculus*), nematode (*Caenorhabditis elegans*), and fruit fly (*Drosophila melanogaster*). Because local alignment is not appropriate for evaluating the overall percentage similarity between protein sequences, we adopted global alignment as an alternative as insects are far distant from well-annotated reference organisms. The program GGSEARCH in the FASTA package [25] was used for implementation of the global alignment search, and organism specific homology searches by GGSEARCH were executed against the four functionally well-annotated organisms described above followed by a search against UniProtKB [34]. A corresponding functional description of the top hit in the reference organism or UniProtKB was annotated as “homolog” (Figure 2).

**Figure 2.**
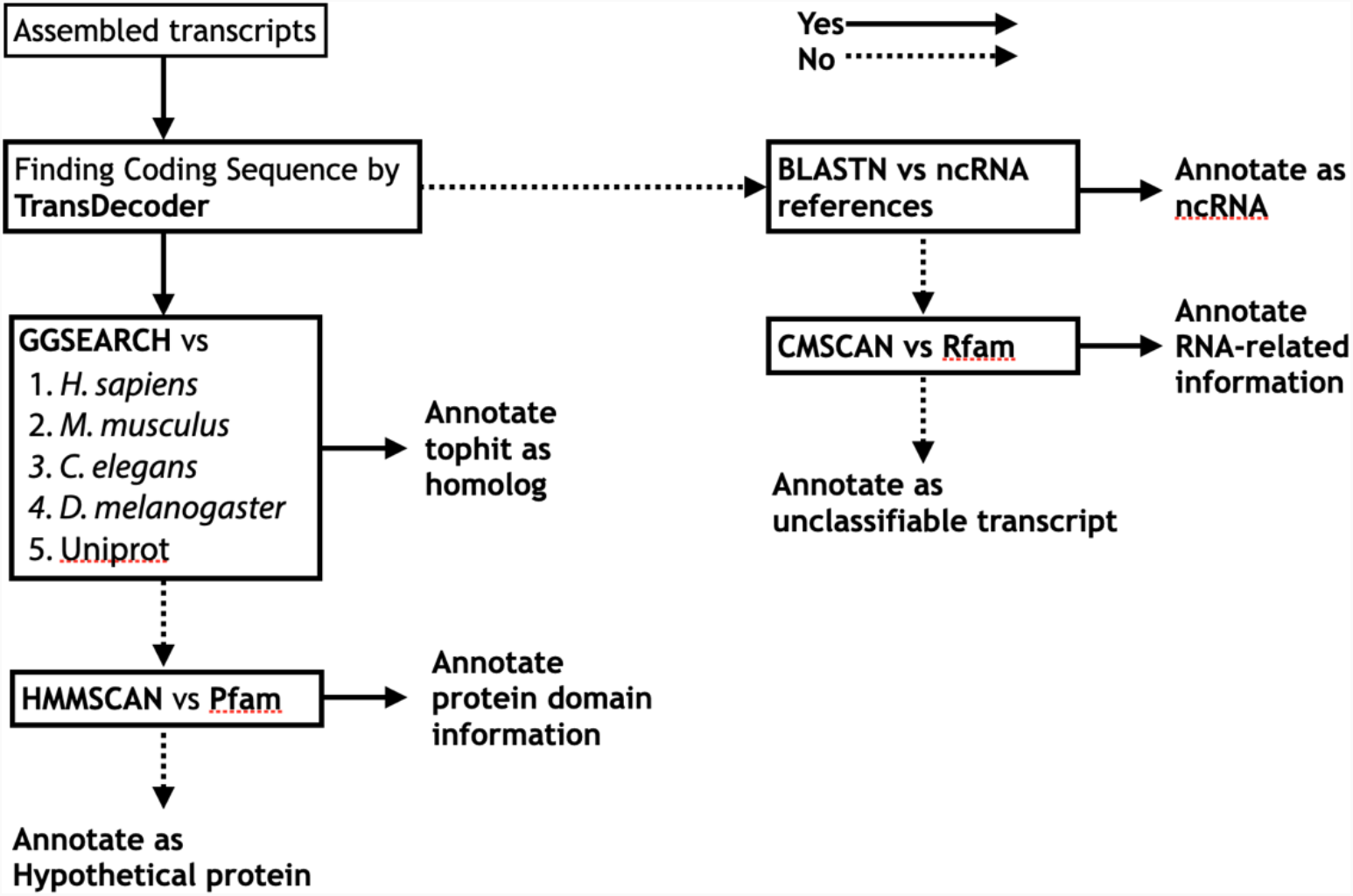
Annotation workflow for transcript description from sequence information. Left half for protein sequence level annotation; right half for nucleotide sequence level annotation.

If no homologs could be found, the search result of HMMSCAN against Pfam was referenced [27] (Figure 2). According to the protein domain hits, protein domain information was annotated if present (example: “zinc finger containing protein”); else, just “hypothetical protein” was used. The sources of the reference DBs used in the functional annotation are summarized in Table 1.

#### 3.1.2. Functional annotation of non-coding sequences

For sequences in which TransDecoder was unable to find any protein coding region (i.e., > 100 amino acids (aa) in length by default), functional information of the ncRNAs were annotated based on nucleotide Basic Local Alignment Search Tool (BLAST) search against ncRNA reference sequences (Figure 2). For insect functional annotation, reference ncRNA sequences of the four functionally well-annotated organisms (*H. sapiens, M. musculus, C. elegans*, and *D. melanogaster*) were used in Fanflow4Insects (Table 1).

If no functional information could be assigned, then the RNA structure and sequence similarity against Rfam [29] were assessed using the CMSCAN program in the Infernal package [28] (Figure 2). If any information regarding the sequence was located, the RNA-related information in DB hits was annotated; else, the sequence was annotated as an “unclassifiable transcript”.

#### 3.1.3. Functional annotation by quantified expression information

Transcript expression was quantified based on the assembled transcriptome and RNA-Seq raw reads. The matrix of gene expression was generated by pasting columns. Using the expression matrix, we were able to annotate gene expression features for all genes.

Although the threshold for the judgment remains to be calibrated, genes with specific expression were then annotated (Figure 3). If a transcript is expressed in all tissues, it was annotated as “constitutive expression”. Alternatively, if expressed only in e.g. the fat body, it was annotated as “fat body-specific expression”. Practical examples of annotation based on expression information are described in the following section.

**Figure 3.**
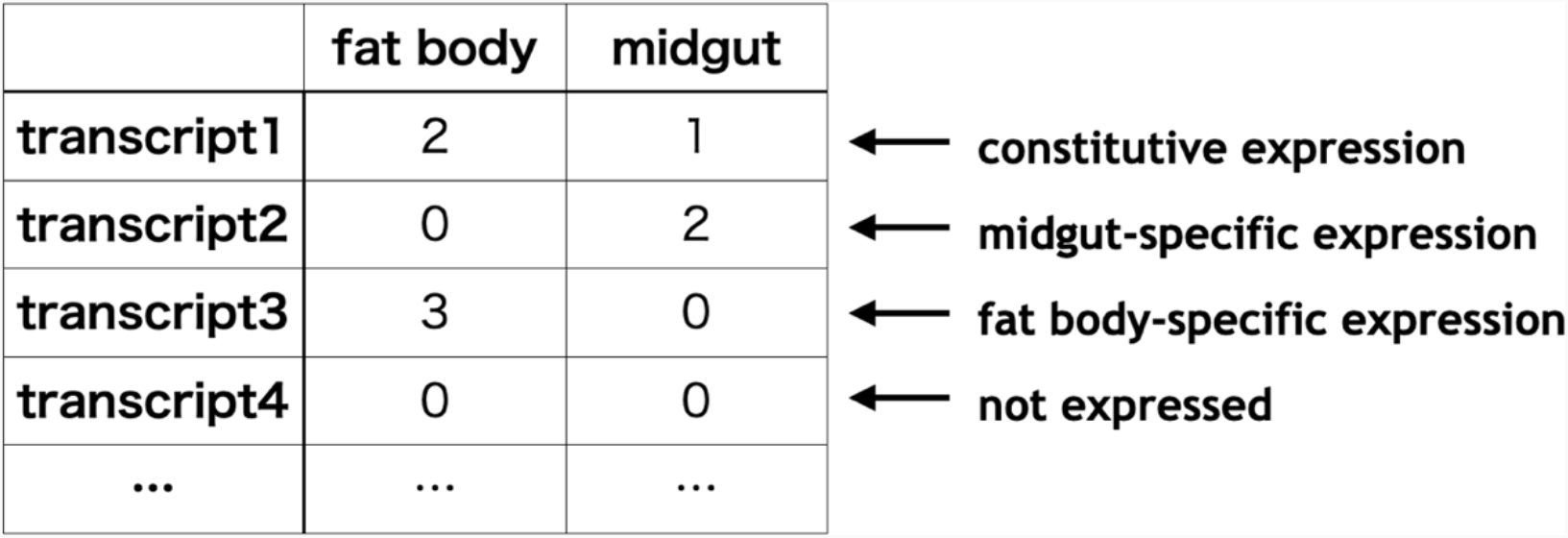
Annotation workflow for transcript description from expression information.

### 3.2. Case study for Fanflow4Insects

Practical applications of Fanflow4Insects for the Japanese stick insect *Entoria okinawaensis* and silkworm *Bombyx mori* are described below.

#### 3.2.1. Functional annotation of the Japanese stick insect transcriptome using Fanflow4Insects

In addition to three biological replicates of *E. okinawaensis* midgut sample, which we previously reported and deposited in SRA under DRA007226 (DRR148118, DRR148119 and DRR148120) [32], three biological replicate reads of *E. okinawaensis* fat body (DRR346171, DRR346172 and DRR346173) were used for the functional annotation by Fanflow4Insects. Transcriptome completeness analysis with BUSCO v5.3.2 using insecta_odb10 dataset (2020-09-10) with default settings (transcriptome mode) showed 97.1% completeness [43.5% single-copy and 53.6% duplicated] for transcriptomes from fat body and midgut of *E. okinawaensis* while it showed 93.0 % completeness [47.9% single-copy and 45.1% duplicated] for that only from midgut of *E. okinawaensis* [35].

Of 311,357 transcripts, 18.5% of those (57,732 transcripts) could be translated into 68,162 predicted protein sequences with more than 100 amino acids length. Some transcripts were predicted to encode more than one protein sequence. These protein coding sequences were annotated by systematic sequence similarity and protein domain searches (Table 2). In brief, 55,068 predicted protein sequences (80.8%) had homologs and 58,876 of those (86.4%) had somewhat sequence similarity information including protein domains. 9,286 predicted protein sequences cannot be assigned to any functional information from sequence similarity and domain searches in protein level and thus annotated as ‘hypothetical protein’.

**Table 2.** Protein-level annotation for *E. okinawaensis*.

| Annotation category | Annotation level | Number of annotation | Cumulative number | Percentage |
| --- | --- | --- | --- | --- |
| Protein homolog from tophit | Human or mouse homolog | 44,351 | 44,351 | 65.1 |
|  | *C. elegans* homolog | 3,739 | 48,090 | 70.6 |
|  | *D. melanogaster* homolog | 2,349 | 50,439 | 74.0 |
|  | Homolog found in UniProtKB | 4,629 | 55,068 | 80.8 |
| No protein homolog | Protein domain | 3,808 | 58,876 | 86.4 |
|  | Hypothetical protein | 9,286 | 68,162 | 100 |

The remaining 253,625 sequences comprised transcripts without protein coding regions. Of these, only 185 sequences had sequence similarity to known ncRNA sequences. Moreover, 264 sequences could be assigned based on a CMSCAN search against Rfam DB, whereas 253,176 sequences were annotated as an “unclassifiable transcript”. These functional annotation results from sequence information for *E. okinawaensis* are summarized graphically in Figure S1.

For the hypothetical protein and the unclassifiable transcripts with no sequence similarity, quantified expression values were used for annotation. Functional information could be inferred and annotated based on the expression matrix generated from sequence counts (Figure 1). Table 3 summarizes the numbers of sequences annotated as “hypothetical protein” and “unclassifiable transcript” and the corresponding expression annotations. Only a small fraction of sequences, 2.7% (156) of “hypothetical proteins” and 1.5% (3,860) of “unclassifiable transcripts”, had no expression information (Table 3).

**Table 3.** Functional annotation from expression for *E. okinawaensis*.

| Annotation level | All (311,357) | Hypothetical protein (5,699) | Unclassifiable transcript (253,176) |
| --- | --- | --- | --- |
| Fat body specific expression | 48,675 | 622 | 42,790 |
| Midgut specific expression | 28,918 | 315 | 25,005 |
| Constitutive expression | 228,103 | 4,606 | 181,521 |
| Not expressed | 5,661 | 156 | 3,860 |

The assembled transcriptomes and the estimated transcript abundances were deposited to the TSA Sequence database under accession IDs: ICSG01000001–ICSG01311357 and the Genomic Expression Archive (GEA) under accession ID: E-GEAD-476. Full functional annotation and transcript descriptions generated by Fanflow4Insects were deposited to figshare to be reused in stick insect research.

#### 3.2.2. Functional annotation of silkworm using Fanflow4Insects

We also evaluated the reference transcriptome data of silkworm *B. mori* as a lepidopteran insect with biological and industrial importance [12]. Corresponding functional annotation is available from the Life Science Database Archive [36]; however, only BLAST top hits for human and fly are listed. We therefore applied Fanflow4Insects to silkworm using the assembled transcriptomes (TSA IDs: ICPK01000001–ICSG01051926) and estimated abundance of transcripts (GEA ID: E-GEAD-315). Transcriptome completeness analysis with BUSCO v5.3.2 for *B. mori* trasncriptome by lepidoptera_odb10 dataset (2020-08-05) yields 98.7% completeness [41.0% single-copy, 57.7% duplicated] [35]. Of 51,926 transcripts, 78.1% (40,555 transcripts) could be translated into 45,719 predicted protein sequences of length > 100 aa. These protein coding sequences were annotated via systematic sequence similarity and protein domain searches, with 38,107 predicted protein sequences (83.4%) able to be annotated with homologs (Table 4). Moreover, protein domains could be assigned to 5,398 predicted protein sequences for the remaining 7,612 predicted protein sequences without homologs, resulting in only 2,214 predicted protein sequences (4.8%) being annotated as a “hypothetical protein” (Table 4).

**Table 4.**
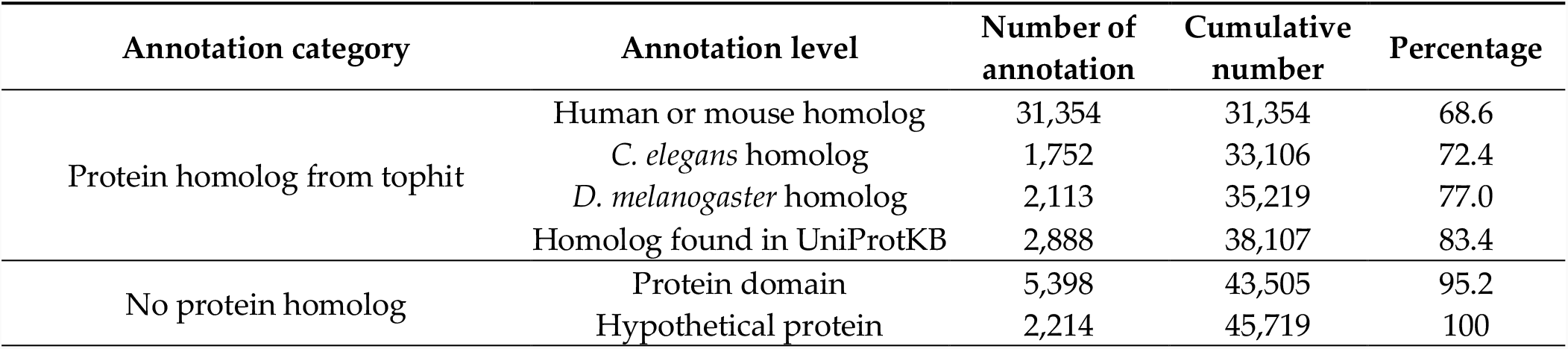
Protein-level annotation for *B. mori*.

The remaining 11,371 sequences comprised transcripts without protein coding regions. Of these, only 45 sequences had sequence similarity to known ncRNA sequenced. Moreover, 300 sequences could be assigned based on CMSCAN search against Rfam DB. Finally, 11,026 sequences were annotated as an “unclassifiable transcript”. These functional annotation results from sequence information for *B. mori* are summarized graphically in Figure S2.

The estimated abundances of transcripts in the expression matrix (GEA ID: E-GEAD-315) were also used for annotation. Table 5 summarizes the numbers of sequences annotated as “hypothetical protein” and “unclassifiable transcript”, together with the corresponding expression annotations. Unlike the results for *E. okinawaensis* in which data from only two tissues were available, the abundance of transcripts in six tissues, notated as transcripts per million (TPM), was available for *B. mori*. A small number of sequences were judged to be tissue-specific because only the genes with the expression in specific tissues (TPM > 0) and no expression (TPM = 0) in other tissues were considered to be tissue specific. In summary, 3.0% (44) of 1,486 transcripts carrying the “hypothetical protein” annotation and 5.4% (693) of 12,845 annotated as an “unclassifiable transcript” could be annotated with strong expression features, although the function of those transcripts could not be inferred based only on sequence similarity information (Table 5).

**Table 5.** Functional annotation of *B.mori* transcripts based on expression.

| Annotation level | All<br>(51,927) | Hypothetical<br>protein<br>(1,486) | Unclassifiable<br>transcript<br>(12,845) |
| --- | --- | --- | --- |
| Fat body-specific expression | 39 | 1 | 11 |
| Midgut-specific expression | 108 | 0 | 29 |
| Malpighian tubule-specific expression | 83 | 3 | 14 |
| Silk gland-specific expression | 365 | 12 | 92 |
| Testis-specific expression | 861 | 19 | 492 |
| Ovary-specific expression | 179 | 9 | 55 |
| Constitutive expression | 7,825 | 114 | 1,205 |
| No expression | 609 | 24 | 125 |

The functional annotation described above including the transcript descriptions generated by Fanflow4Insects along with the expression annotation has been deposited to figshare for use in silkworm research.

#### 3.2.3. Comparison of ncRNAs between Japanese stick insect and silkworm

A comparative analysis of ncRNA transcripts was performed to utilize the output of Fanflow4Insects. Notably, the functional annotation to ncRNA references (human and fruit fly) by Fanflow4Insects provided transcript IDs for *E. okinawaensis* and *B. mori* along with corresponding ncRNA IDs from reference organisms for use in this comparison (Figure 4).

**Figure 4.**
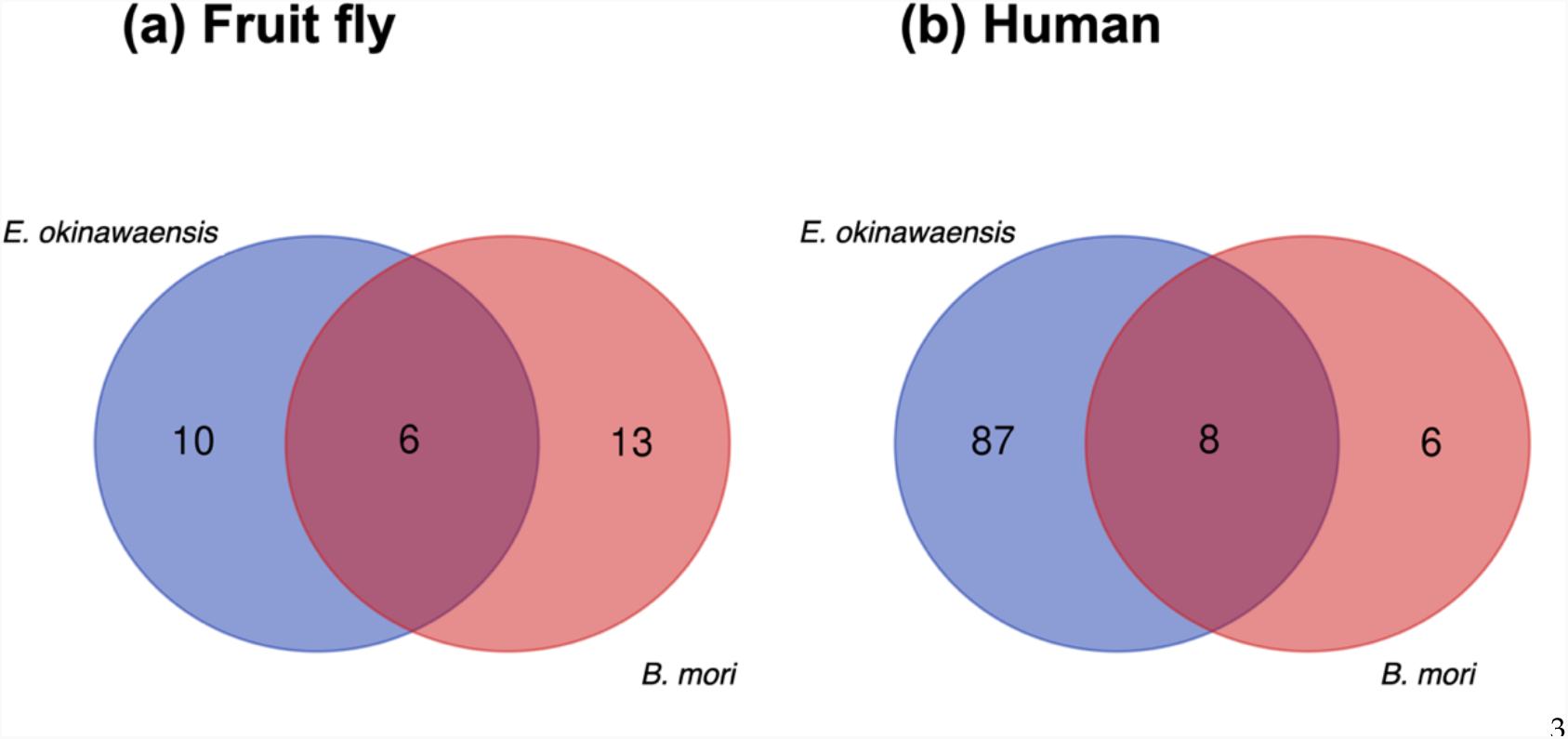
Comparison of ncRNA transcripts between *E. okinawaensis* and *B. mori* (a) The numbers of ncRNA transcripts predicted from fruit fly annotation. (b) The numbers of ncRNA transcripts predicted from human annotation.

Whereas few transcripts were functionally annotated based on existing ncRNAs in the public database, several functionally annotated transcripts were identified in the intersection of *E. okinawaensis* and *B. mori*; the corresponding transcript IDs in the reference organisms (human and fruit fly) are listed in Table 6. All *E. okinawaensis* transcripts annotated and then listed in Table 6 have more than one counterpart transcripts in *B. mori* by BLASTN search. We also evaluated the corresponding functional annotation derived from the expression of these transcripts, revealing that most transcripts exhibit constitutive expression whereas several show midgut-specific expression in *E. okinawaensis*; in contrast, all transcripts are constitutively expressed in *B. mori*.

**Table 6.**
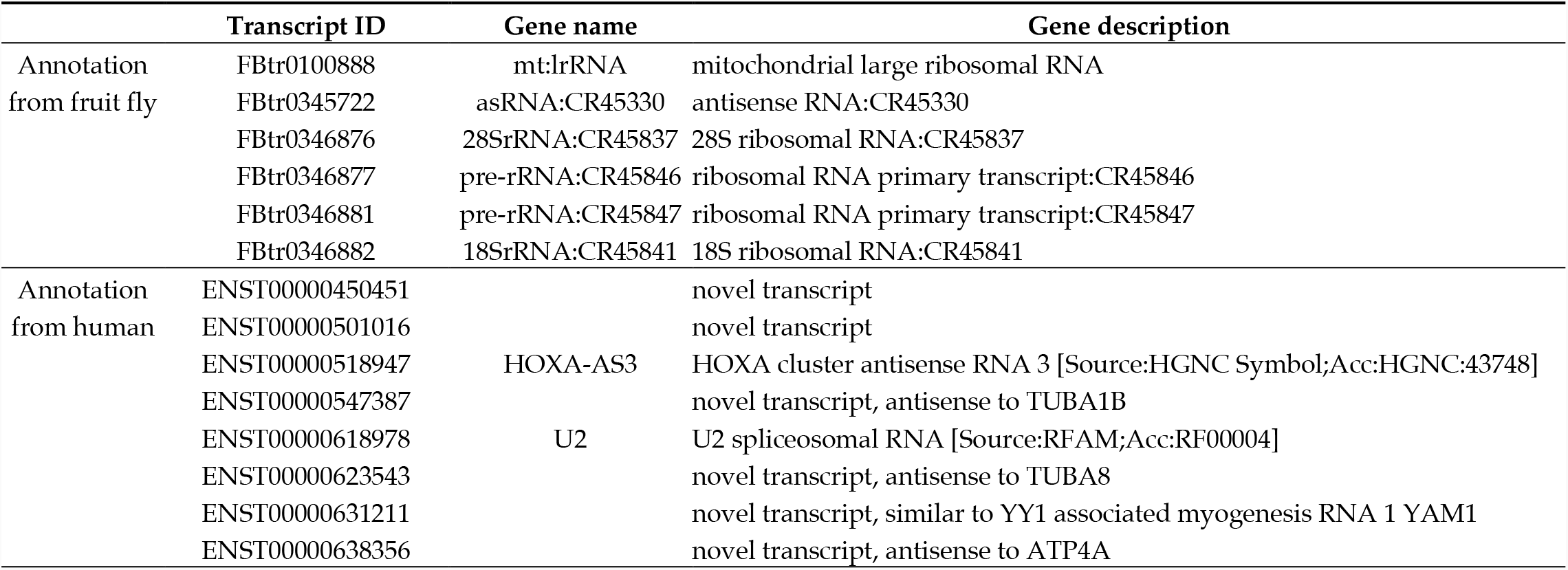
ncRNA transcripts expressed both in *E. okinawaensis* and *B. mori*.

## 4. Discussion

Next-generation sequencing has facilitated decoding of the genomes of numerous insects along with their transcript sequences. However, the biological interpretation of these sequences remains as a primary bottleneck of transcriptome analysis. An important first step is functional annotation, which serves as e.g. an important clue for selecting genome editing targets. Toward this end, we have had a long-term interest in providing functional annotation, starting with the annotation of enzymatic genes using the Gene Function Identification Tool (GFIT) in bacteria [37]. More recently, we have contributed to the functional annotation pipeline in the FANTOM project to provide the scientific community with accurate functional annotation of mouse cDNA [15-17].

Currently, BLAST2GO is often used for functional annotation of non-model organisms [4]; however, BLAST2GO is a commercial application and considerable delay exists before results are obtained. Recently, Seq2fun has been used to predict the function of the total set of genes encoded in a genome at very high speed [5]. However, Seq2fun does not generate sequences for each gene, making it unusable for applications such as individual transcriptome analysis. Thus, it is not possible to trace how the functional annotation was assigned to each gene using Seq2fun. We consider that traceable functional annotation supported by concrete evidence is crucial in biological studies. In addition, these tools cannot be customized to the circumstances of each organism. To address these issues, we therefore developed Fanflow4Insects.

In our previous study for annotating the *E. okinawaensis* midgut transcriptome, we performed a systematic sequence similarity search using translated protein sequences against the predicted protein sequences of well-annotated model organisms using a locally installed NCBI protein BLAST program [32], which provided sufficient speeds to obtain results in a reasonable time. However, although BLAST reports the local alignment of the query sequence and DB hit sequence [28], the local alignment is not appropriate for the present purposes because overall sequence identity and sequence similarity between protein sequences cannot be determined. Therefore, in the present study we re-evaluated the program to be applied for the project. The program FASTY in the FASTA package [25], previously utilized in the FANTOM project [14, 18], compares a DNA sequence to a protein sequence DB, translating the DNA sequence in three forward (or reverse) frames and allowing frameshifts. However, despite dramatic improvements in computation technology, FASTY still requires considerable computing power and is difficult to be used for practical functional annotation. Considering that next-generation sequencing technology using “sequence by synthesis” with Illumina sequencers is much more accurate as compared to capillary sequencing using the Sanger method, and nucleotide deletions are less likely to occur in the final sequences, we proposed to utilize global rather than local alignment in the present study. Global alignment is also more appropriate because several well-conserved alignment fragments are apparent upon the alignment of insects and model organisms while insects are far distant from the well-annotated reference organisms. We found that the GGSEARCH program in the FASTA package is most suitable for the implementation of global alignment searches in a local installation environment. In particular, GGSEARCH calculates sequence identity and similarity, which allow biologists to intuitively evaluate the homology between two sequences, using global alignment while providing similar speed to BLAST. However, because of the evolutionary distance between insects and well-annotated model organisms, we set a lower E-value threshold of 0.1 in the systematic sequence similarity search by GGSEARCH against genes from well-annotated organisms (human, mouse, *C. elegans*, and *D. melanogaster*) followed by UniProtKB [34] (Figure 2). By this procedure, the functional annotation of genes was dramatically improved by using well-annotated species protein sequence datasets compared with the use of only non-redundant dataset (nr), where so many genes tend to be annotated as ‘hypothetical protein’ because gene descriptions of related organisms are frequently assigned. The use of silkworms and Japanese stick insects as well as parasitic wasps *Copidosoma floridanum* made this feature suitable for functional annotation of insect species possible [38,39].

In practice, more than 80% of protein sequences for *E. okinawaensis* and *B. mori* could be assigned homologs using GGSEARCH (Table 2 and 4). In our previous study, we showed that approximately 58% of *B. mori* genes had human homologs [11]. In the present study, we showed that 68.6% of *B. mori* protein sequences could be assigned human or mouse homologs (Table 4). We believe that the increase in percentage directly reflects advances in our knowledge of reference sequences.

Although many DBs have been developed for protein domains, Pfam [27], which describes protein domains using the Hidden Markov Model (HMM), is often used. We therefore included the program HMMSCAN in the HMMER package [26], available as a program to search Pfam, in the workflow. Notably, we obtained functional information for numerous sequences for which no sequence similarity to the reference protein sequence was identified using protein domain assignment. In particular, some protein domain information could be assigned for 3,808 sequences in *E. okinawaensis* and 5,398 sequences in *B. mori* (Table 2 and 4), which allows the function of the sequence to be inferred despite the lack of similarity to a known protein sequence. Overall, this strategy allowed 95.2% of protein sequences in *B. mori* to be annotated with only 4.8% (2,214 sequences) being left unannotated. These findings indicate that sequence similarity and protein domain searches with proper reference databases (Table 1) can serve as a powerful tool for functional annotation.

The accumulation of ncRNA reference sequences in model organisms facilitates sequence-level comparisons; thus, we incorporated ncRNAs incorporated into the workflow. Toward this end, we utilized nucleotide BLAST, as sequences are not as conserved at the nucleotide sequence level as at the protein sequence level. Therefore, although few assignments were obtained, they were useful to infer functionality. In particular, knowledge regarding RNA domains has been accumulated in a DB called Rfam [30], with RNA domains able to be searched using the program CMSCAN in the Infernal package [29]; thus, this program was also incorporated into the workflow. We anticipate that although only a few sequences were annotated from ncRNA DB and Rfam for *E. okinawaensis* and *B. mori* (Figures S1 and S2), additional sequences will be annotated in the future as ncRNA reference sequences and RNA domain information become more complete. While we utilized CMSCAN program to search against Rfam DB in the current version of Fanflow4Insects, the functional annotation of ncRNA should be improved in a coming version of Fanflow4Insects to draw functional inference from existing databases. While the current target of Fanflow4Insects is to annotate functional information from public databases for typical RNAseq reads of insect species, we should be careful about the information of RNAseq protocols in the annotation of ncRNAs.

Performing functional annotation based on gene expression is challenging, encompassing numerous problems such as how to determine the threshold and the inability to detect weak expression unless the RNA-Seq reads are sufficiently deep [40]. However, gene expression can provide useful information for inferring the function of the sequence. Toward this end, information from derived libraries has been previously used in the analysis of ESTs [15–17]. More recently such information has been replaced by quantified expression values derived from RNA-Seq, which can be used for functional annotation together with the level of expression in other tissues. Accordingly, we also integrated quantified expression values for providing functional annotation. In the present study, sequences that were expressed only in a specific tissue were annotated as showing tissue-specific expression (Table 3 and 5). Notably, as quantified expression data were available for only two tissues in *E. okinawaensis*, multiple sequences were judged as tissue specific. This allowed us to add functional annotation based on expression to sequences previously annotated as “hypothetical protein” or “unclassifiable transcript” (Table 3). In comparison, as data for six tissues were available in *B. mori* only few sequences exhibited tissue specific expression and could accordingly be assigned functional annotation (Table 5). Moreover, the functional annotation to ncRNA references in Fanflow4Insects allowed the comparative analysis of ncRNA transcripts (Figure 4). In conjunction with the expression-based functional annotation, the comparative use of Fanflow4Insects data could therefore readily identify candidate genes for future evaluation of e.g. insects with distinct phenotypes (Table 6). These results indicated that even sequences for which no information was obtained from sequence similarity were able to provide useful functional information.

For enrichment analysis, it is especially important to render the data not only human-readable with regard to transcript description but also machine-readable using GO and other functional annotations. Although originally developed by fly geneticists, GO is now widely used in various model organisms [7]. In our previous study for annotating the *E. okinawaensis* midgut transcriptome, enrichment analyses were performed by assigning transcripts of *E. okinawaensis* to those of *D. melanogaster* because no original GO annotations were available for *E. okinawaensis* [32]. The next target is to annotate GO terms to newly sequenced organism and utilize those for enrichment analysis. Currently, GO annotation applies only for protein coding sequences; however, we speculate that annotation using a controlled vocabulary including that of GO will be incorporated for functional annotation of both proteins and transcripts because it affords machine-readability. As the genome sequences of more organisms become available, functional annotation of the genes encoded therein will be required together with GO and protein domain annotation to e.g. play an active role in selecting which genes to edit for genome editing.

Following the Findable, Accessible, Interoperable, Re-usable (FAIR) principle [41], raw RNA-Seq data and hopefully assembled transcriptome sequences should be registered in the public DB as well as to reproduce data analysis. Our data regarding RNA-Seq reads, transcriptome assembly, and quantified expression values are archived in the public DBs Sequence Read Archive (SRA), TSA, and GEA, respectively. A workflow that integrates these elements is very important and therefore should be re-usable with proper interoperability. Developed openly on GitHub, so it can be easily cloned from GitHub repositories. Also, specific tools including GGSEARCH can easily be introduced through bioconda framework even if users are not a bioinformatician [42]. All scripts to run the program described in the present study are available on Github: https://github.com/bonohu/SAQE/. The program is not fully automated as it is expected to be customized for actual use. In particular, gene expression information will vary with each insect species and data acquisition situation. Moreover, the calculation of correspondence relationships with closely related insects for which sequence information is already available will also vary depending on the situation. Such construction of individual workflows according to the available information represents the key to digital transformation while brand-new function in target insects cannot be assigned.

## 5. Conclusions

We have developed a functional annotation workflow for insects, termed “Fanflow4Insects”. Fanflow4Insects has been openly developed on GitHub, and is freely accessible. Fanflow4Insects was tested for functional annotation of the Japanese stick insect *E. okinawaensis* and silkworm *B. mori*. In conjunction with functional annotation derived from expression, the data from Fanflow4Insects can be applicable to the comparative study of insects with distinct phenotypes.

## Supplementary Materials

Figure S1: Graphical summary of functional annotation from sequence information for *E. okinawaensis*. Figure S2: Graphical summary of functional annotation from sequence information for *B. mori*.

## Author Contributions

Conceptualization, H.B. and T.K.; methodology, H.B.; software, H.B.; validation, H.B. and T.K.; formal analysis, H.B.; investigation, T.S and H.B.; resources, T.S, H.T. and H.B.; data curation, H.B.; writing—original draft preparation, H.B.; writing—review and editing, H.B., T.S., H.T. and T.K.; visualization, H.B.; supervision, H.B.; project administration, H.B.; funding acquisition, H.B. and H.T. All authors have read and agreed to the published version of the manuscript.

## Funding

This research was supported by JSPS KAKENHI Grant 21K19118 to HB. This work was also supported by the Center of Innovation for Bio-Digital Transformation (BioDX), an open innovation platform for industry-academia co-creation (COI-NEXT), Japan Science and Technology Agency (JST, COI-NEXT, JPMJPF2010) to HB and the ROIS-DS-JOINT (004RP2017) to HT. The APC was funded by JSPS KAKENHI Grant 21K19118.

## Institutional Review Board Statement

Not applicables.

## Data Availability Statement

Workflow developed and original scripts to replicate the study are available on GitHub: https://github.com/bonohu/SAQE/. The RNA sequencing reads reported in this article are available in the Sequence Read Archive (SRA) under the accession DRA013458 (DRR346171, DRR346172 and DRR346173). The assembled transcriptome sequences are available in the Transcriptome Shotgun Assembly (TSA) sequence database under the accession ICSG01(ICSG01000001-ICSG01311357), and the estimated abundance of transcripts is available from the Genomic Expression Archive (GEA) under the accession ID E-GEAD-476.

## Acknowledgments

Authors wish to thank Dr. Kakeru Yokoi for helpful discussion. Computations were performed on the computers at Hiroshima University Genome Editing Innovation Center.

## Conflicts of Interest

The authors declare no conflict of interest. The funders had no role in the design of the study; in the collection, analyses, or interpretation of data; in the writing of the manuscript, or in the decision to publish the results.

## Notes

https://doi.org/10.6084/m9.figshare.19368110

https://doi.org/10.6084/m9.figshare.19368137

### Competing Interest Statement

The authors have declared no competing interest.

### Summary of Updates

Discussion added.

## References

1. Genomes – NCBI Datasets for Insecta; doi:10.1093/gbe/evab138. https://www.ncbi.nlm.nih.gov/datasets/genomes/?taxon=50557 Accessed 14 May 2022.

2. Hotaling, S.; Sproul, J.S.; Heckenhauer, J.; Powell, A.; Larracuente, A.M.; Pauls, S.U.; Kelley, J.L.; Frandsen, P.B. Long Reads Are Revolutionizing 20 Years of Insect Genome Sequencing. Genome Biol Evol 2021, 13, evab138, doi:10.1093/gbe/evab138.

3. Oppenheim, S.J.; Baker, R.H.; Simon, S.; DeSalle, R. We Can’t All Be Supermodels: The Value of Comparative Transcriptomics to the Study of Non-Model Insects. Insect Mol Biol 2015, 24, 139–154, doi:10.1111/imb.12154.

4. Conesa, A.; Götz, S.; García-Gómez, J.M.; Terol, J.; Talón, M.; Robles, M. Blast2GO: A Universal Tool for Annotation, Visualization and Analysis in Functional Genomics Research. Bioinformatics 2005, 21, 3674–3676, doi:10.1093/bioinformatics/bti610.

5. Liu, P.; Ewald, J.; Galvez, J.H.; Head, J.; Crump, D.; Bourque, G.; Basu, N.; Xia, J. Ultrafast Functional Profiling of RNA-Seq Data for Nonmodel Organisms. Genome Res 2021, 31, 713–720, doi:10.1101/gr.269894.120.

6. Larkin, A.; Marygold, S.J.; Antonazzo, G.; Attrill, H.; Dos Santos, G.; Garapati, P.V.; Goodman, J.L.; Gramates, L.S.; Millburn, G.; Strelets, V.B.; et al. FlyBase: Updates to the Drosophila Melanogaster Knowledge Base. Nucleic Acids Res 2021, 49, D899– D907, doi:10.1093/nar/gkaa1026.

7. Ashburner, M.; Ball, C.A.; Blake, J.A.; Botstein, D.; Butler, H.; Cherry, J.M.; Davis, A.P.; Dolinski, K.; Dwight, S.S.; Eppig, J.T.; et al. Gene Ontology: Tool for the Unification of Biology. The Gene Ontology Consortium. Nat Genet 2000, 25, 25–29, doi:10.1038/75556.

8. Cunningham, F.; Allen, J.E.; Allen, J.; Alvarez-Jarreta, J.; Amode, M.R.; Armean, I.M.; Austine-Orimoloye, O.; Azov, A.G.; Barnes, I.; Bennett, R.; et al. Ensembl 2022. Nucleic Acids Res 2022, 50, D988–D995, doi:10.1093/nar/gkab1049.

9. Yates, A.D.; Allen, J.; Amode, R.M.; Azov, A.G.; Barba, M.; Becerra, A.; Bhai, J.; Campbell, L.I.; Carbajo Martinez, M.; Chakiachvili, M.; et al. Ensembl Genomes 2022: An Expanding Genome Resource for Non-Vertebrates. Nucleic Acids Res 2022, 50, D996– D1003, doi:10.1093/nar/gkab1007.

10. Tabunoki, H.; Ono, H.; Ode, H.; Ishikawa, K.; Kawana, N.; Banno, Y.; Shimada, T.; Nakamura, Y.; Yamamoto, K.; Satoh, J.-I.; et al. Identification of Key Uric Acid Synthesis Pathway in a Unique Mutant Silkworm Bombyx Mori Model of Parkinson’s Disease. PLoS One 2013, 8, e69130, doi:10.1371/journal.pone.0069130.

11. Tabunoki, H.; Bono, H.; Ito, K.; Yokoyama, T. Can the Silkworm (Bombyx Mori) Be Used as a Human Disease Model? Drug Discov Ther 2016, 10, 3–8, doi:10.5582/ddt.2016.01011.

12. Yokoi, K.; Tsubota, T.; Jouraku, A.; Sezutsu, H.; Bono, H. Reference Transcriptome Data in Silkworm Bombyx Mori. Insects 2021, 12, 519, doi:10.3390/insects12060519.

13. Nojima, Y.; Bono, H.; Yokoyama, T.; Iwabuchi, K.; Sato, R.; Arai, K.; Tabunoki, H. Superoxide Dismutase Down-Regulation and the Oxidative Stress Is Required to Initiate Pupation in Bombyx Mori. Sci Rep 2019, 9, 14693, doi:10.1038/s41598-019-51163-3.

14. Kasukawa, T.; Furuno, M.; Nikaido, I.; Bono, H.; Hume, D.A.; Bult, C.; Hill, D.P.; Baldarelli, R.; Gough, J.; Kanapin, A.; et al. Development and Evaluation of an Automated Annotation Pipeline and CDNA Annotation System. Genome Res 2003, 13, 1542– 1551, doi:10.1101/gr.992803.

15. Kawai, J.; Shinagawa, A.; Shibata, K.; Yoshino, M.; Itoh, M.; Ishii, Y.; Arakawa, T.; Hara, A.; Fukunishi, Y.; Konno, H.; et al. Functional Annotation of a Full-Length Mouse CDNA Collection. Nature 2001, 409, 685–690, doi:10.1038/35055500.

16. Okazaki Y, Furuno M, Kasukawa T, Adachi J, Bono H, Kondo S, et al. Analysis of the mouse transcriptome based on functional annotation of 60,770 full-length cDNAs. Nature. 2002;420:563–73; doi:10.1038/nature01266.

17. Carninci, P.; Kasukawa, T.; Katayama, S.; Gough, J.; Frith, M.C.; Maeda, N.; Oyama, R.; Ravasi, T.; Lenhard, B.; Wells, C.; et al. The Transcriptional Landscape of the Mammalian Genome. Science 2005, 309, 1559–1563, doi:10.1126/science.1112014.

18. Maeda, N.; Kasukawa, T.; Oyama, R.; Gough, J.; Frith, M.; Engström, P.G.; Lenhard, B.; Aturaliya, R.N.; Batalov, S.; Beisel, K.W.; et al. Transcript Annotation in FANTOM3: Mouse Gene Catalog Based on Physical CDNAs. PLoS Genet 2006, 2, e62, doi:10.1371/journal.pgen.0020062.

19. Blaxter, M.; Danchin, A.; Savakis, B.; Fukami-Kobayashi, K.; Kurokawa, K.; Sugano, S.; Roberts, R.J.; Salzberg, S.L.; Wu, C.-I. Reminder to Deposit DNA Sequences. Science 2016, 352, 780, doi:10.1126/science.aaf7672.

20. Salzberg, S.L. Databases: Reminder to Deposit DNA Sequences. Nature 2016, 533, 179, doi:10.1038/533179a.

21. Bono H. Meta-analysis of oxidative transcriptomes in insects. Antioxidants (Basel). 2021, 10:345; doi:10.3390/antiox10030345.

22. Github repository: bonohu/SAQE. https://github.com/bonohu/SAQE Accessed on 14 May 2022.

23. Grabherr, M.G.; Haas, B.J.; Yassour, M.; Levin, J.Z.; Thompson, D.A.; Amit, I.; Adiconis, X.; Fan, L.; Raychowdhury, R.; Zeng, Q.; et al. Full-Length Transcriptome Assembly from RNA-Seq Data without a Reference Genome. Nat Biotechnol 2011, 29, 644– 652, doi:10.1038/nbt.1883.

24. Haas, B.J.; Papanicolaou, A.; Yassour, M.; Grabherr, M.; Blood, P.D.; Bowden, J.; Couger, M.B.; Eccles, D.; Li, B.; Lieber, M.; et al. De Novo Transcript Sequence Reconstruction from RNA-Seq Using the Trinity Platform for Reference Generation and Analysis. Nat Protoc 2013, 8, 1494–1512, doi:10.1038/nprot.2013.084.

25. FASTA Sequence Comparison at the U. of Virginia. https://fasta.bioch.virginia.edu/ Accessed 14 May 2022.

26. HMMER: biosequence analysis using profile hidden Markov models. https://hmmer.org/ Accessed 14 May 2022.

27. Mistry, J.; Chuguransky, S.; Williams, L.; Qureshi, M.; Salazar, G.A.; Sonnhammer, E.L.L.; Tosatto, S.C.E.; Paladin, L.; Raj, S.; Richardson, L.J.; et al. Pfam: The Protein Families Database in 2021. Nucleic Acids Res 2021, 49, D412–D419, doi:10.1093/nar/gkaa913.

28. Altschul, S.F.; Madden, T.L.; Schäffer, A.A.; Zhang, J.; Zhang, Z.; Miller, W.; Lipman, D.J. Gapped BLAST and PSI-BLAST: A New Generation of Protein Database Search Programs. Nucleic Acids Res 1997, 25, 3389–3402, doi:10.1093/nar/25.17.3389.

29. Nawrocki, E.P.; Eddy, S.R. Infernal 1.1: 100-Fold Faster RNA Homology Searches. Bioinformatics 2013, 29, 2933–2935, doi:10.1093/bioinformatics/btt509.

30. Kalvari, I.; Nawrocki, E.P.; Ontiveros-Palacios, N.; Argasinska, J.; Lamkiewicz, K.; Marz, M.; Griffiths-Jones, S.; Toffano-Nioche, C.; Gautheret, D.; Weinberg, Z.; et al. Rfam 14: Expanded Coverage of Metagenomic, Viral and MicroRNA Families. Nucleic Acids Res 2021, 49, D192–D200, doi:10.1093/nar/gkaa1047.

31. Patro, R.; Duggal, G.; Love, M.I.; Irizarry, R.A.; Kingsford, C. Salmon Provides Fast and Bias-Aware Quantification of Transcript Expression. Nat Methods 2017, 14, 417–419, doi:10.1038/nmeth.4197.

32. Sakamoto, T.; Sasaki, S.; Yamaguchi, N.; Nakano, M.; Sato, H.; Iwabuchi, K.; Tabunoki, H.; Simpson, R.J.; Bono, H. De Novo Transcriptome Analysis for Examination of the Nutrition Metabolic System Related to the Evolutionary Process through Which Stick Insects Gain the Ability of Flight (Phasmatodea). BMC Res Notes 2021, 14, 182, doi:10.1186/s13104-021-05600-0.

33. Github repository: FelixKrueger/TrimGalore. https://github.com/FelixKrueger/TrimGalore Accessed on 14 May 2022.

34. UniProt Consortium UniProt: The Universal Protein Knowledgebase in 2021. Nucleic Acids Res 2021, 49, D480–D489, doi:10.1093/nar/gkaa1100.

35. Manni, M.; Berkeley, MR.; Seppey, M.; Simão, FA.; Zdobnov, EM. BUSCO Update: Novel and streamlined workflows along with broader and deeper phylogenetic coverage for scoring of eukaryotic, prokaryotic, and viral genomes. Mol Biol Evol 2021, 38:4647–4654, doi: 10.1093/molbev/msab199.

36. Silkworm, Bombyx mori, reference transcriptome data. https://doi.org/10.18908/lsdba.nbdc02443-002 Accessed 14 May 2022

37. Bono, H.; Ogata, H.; Goto, S.; Kanehisa, M. Reconstruction of Amino Acid Biosynthesis Pathways from the Complete Genome Sequence. Genome Res 1998, 8, 203–210, doi:10.1101/gr.8.3.203.

38. Ohno, H.; Sakamoto, T.; Okochi, R.; Nishiko, M.; Sasaki, S.; Bono, H.; et al. Apoptosis-mediated vasa down-regulation controls developmental transformation in Japanese Copidosoma floridanum female soldiers. Dev Biol 2019, 456, 226–233, doi: 10.1016/j.ydbio.2019.09.005.

39. Sakamoto, T.; Nishiko, M.; Bono, H.; Nakazato, T.; Yoshimura, J.; Tabunoki, H.; Iwabuchi, K. Analysis of molecular mechanism for acceleration of polyembryony using gene functional annotation pipeline in Copidosoma floridanum. BMC Genomics 2020 21, 152, doi: 10.1186/s12864-020-6559-3.

40. Munro SA, Lund SP, Pine PS, Binder H, Clevert D, Conesa A, et al. Assessing technical performance in differential gene expression experiments with external spike-in RNA control ratio mixtures. Nat. Comms. 2014; 5:5125.

41. Wilkinson, M.D.; Dumontier, M.; Aalbersberg, I.J.J.; Appleton, G.; Axton, M.; Baak, A.; Blomberg, N.; Boiten, J.-W.; da Silva Santos, L.B.; Bourne, P.E.; et al. The FAIR Guiding Principles for Scientific Data Management and Stewardship. Sci Data 2016, 3, 160018, doi:10.1038/sdata.2016.18

42. Bioconda. https://bioconda.github.io Accessed 11 June 2022.

