## Supplementary figures and images for "Systematic functional annotation workflow for insects"

### Supplemental Figures

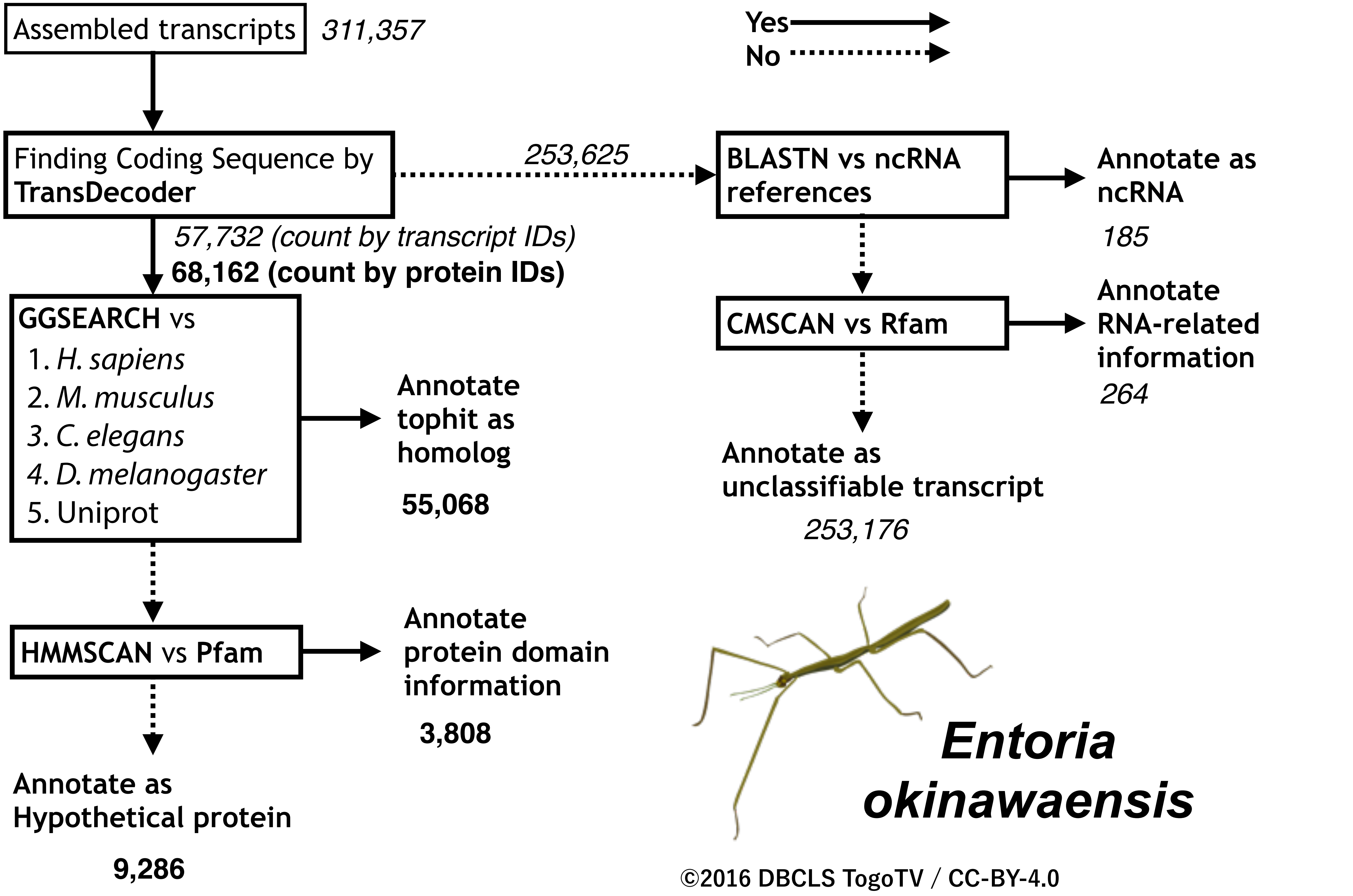

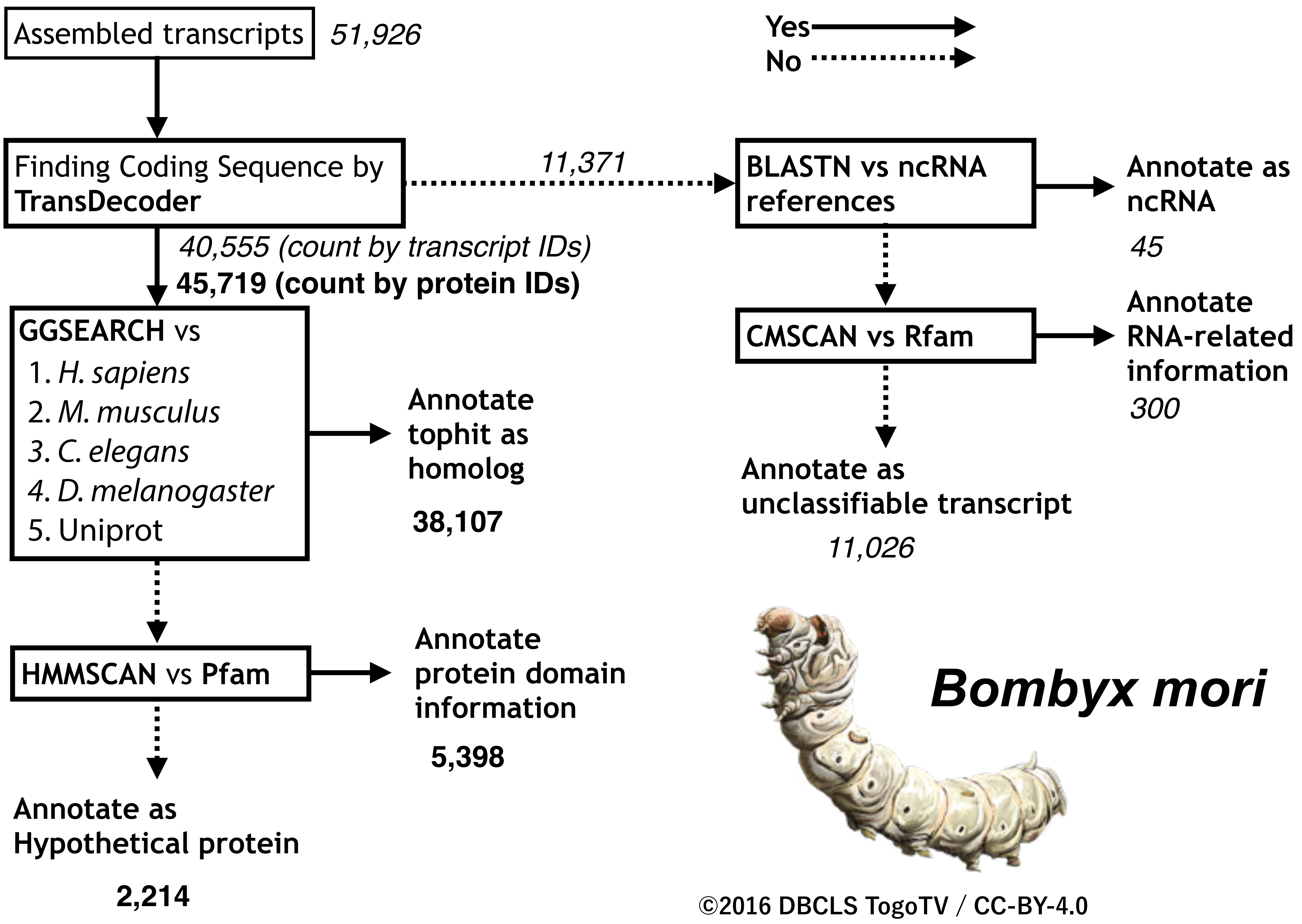
